# The role of microglia membrane potential in chemotaxis

**DOI:** 10.1101/2020.05.19.104109

**Authors:** Laura Laprell, Christian Schulze, Marie-Luise Brehme, Thomas G. Oertner

## Abstract

Microglia react to danger signals by rapid and targeted extension of cellular processes towards the source of the signal. This positive chemotactic response is accompanied by a hyperpolarization of the microglia membrane. Here we show that optogenetic depolarization of microglia has little effect on baseline motility, but significantly slows down the chemotactic response. Reducing the extracellular Ca^2+^ concentration mimics the effect of optogenetic depolarization. As the membrane potential sets the driving force for Ca^2+^ entry, hyperpolarization is an integral part of rapid stimulus-response coupling in microglia. Compared to typical excitable cells such as neurons, the sign of the activating response is inverted in microglia, leading to inhibition by depolarizing channelrhodopsins.

## Background

Microglia – the only resident immune cells in the brain – are the first responders to external and internal threats in the central nervous system. They account for 5-10% of total brain cells and constantly surveil the brain parenchyma. Under physiological conditions microglia are highly ramified, have a small cell body and span approximately 50 µm in diameter. With their thin processes, microglia detect changes in the extracellular space, such as dying neurons, danger signals (e.g. adenine nucleotides), invading organisms, or alterations in neuronal activity [1]. Under physiological conditions, the tandem-pore domain halothane-inhibited K^+^ channel 1 (THIK-1) and possibly transient receptor potential (TRP) and chloride-conducting channels contribute to the resting membrane potential of microglia, typically about - 40 mV [2–4]. Deviations from this potential in both directions have been linked to microglia activation. Under pathological conditions, microglia exhibit two modes on how they change their membrane potential, (i) sustained hyperpolarization in chronically activated microglia and (ii) rapid, transient hyperpolarization induced by extracellular ATP in the vicinity of tissue damage. Sustained pathological hyperpolarization in activated and aged microglia is correlated with an increased expression of Kir2.1 and Kv1.3 channels [4,5]. This additional hyperpolarization is thought to be a precondition for the initiation of microglia functions such as phagocytosis, chemotaxis and cytokine release, which are potentially promoted by the larger driving force for Ca^2+^ influx [6–8]. While sustained hyperpolarization is accompanied by process retraction leading to an ameboid microglia morphology, transient hyperpolarization due to local tissue damage is associated with rapid process extension towards the source of extracellular ATP and Ca^2+^ transients in the processes directed towards the damage site [1,9– 11]. Detection of extracellular ATP and chemotaxis are largely mediated by the purinergic Gi-protein coupled receptor P2Y12 and ion channels receptors P2X4 and P2X7 [12–15].

Although classified as non-excitable cells, microglia exhibit strong electrophysiological stimulus-response features. While correlations between microglia activation and membrane potential are numerous, they have only been described phenomenologically, and it is not well understood whether changes in membrane potential are cause or consequence of microglia activation. Selective investigation of microglia in their native environment is hampered by the fact that identical molecular pathways exist in other types of glial cells and in neurons [4,5,7,16]. Therefore, pharmacological manipulations are extremely difficult to interpret unless microglia are cultured in isolation. In isolation, however, microglia assume an activated phenotype with retracted processes, making it impossible to investigate the activation process from the resting state. Genetic manipulations have the required cell-type specificity, but knockdown or knockout approaches change the system in a chronic fashion and may trigger the activation of compensatory mechanisms. Here we adopt an optogenetic approach to investigate context-dependent behavior of microglia. Using a specific Cre-driver line, we express the light-gated cation channel ChETA specifically in microglia. Two-photon time-lapse imaging allowed us to quantify chemotactic responses while manipulating the membrane potential with blue light pulses. We demonstrate that ATP-mediated hyperpolarization is not an epiphenomenon, but causally related to the chemotactic response.

## Materials and methods

### Mouse slice culture

Mice carrying a tamoxifen-inducible Cre-recombinase in microglia and the floxed fluorescent marker tdTomato (B6.129; B6.129 - Cx3cr1^tm2.1(cre/ERT2)Jung^ Gt(ROSA26)Sortm^9(CAG-tdTomato)Hze^) (JAX 020940; JAX 007909) were crossed with mice heterozygously carrying the Gt(ROSA26)Sor^tm1(CAG-COP4*E123T*H134R,- tdTomato)Gfng^ (JAX 017455) allele to generate mice expressing ChETA and tdTomato in microglia and littermate controls without ChETA expression in microglia. Hippocampal slice cultures from sex-matched ChETA-expressing mice and littermate controls of were prepared at postnatal day 4–7 as described [17]. Briefly, mice were anesthetized with 80% CO_2_ 20% O_2_ and decapitated. Hippocampi were dissected in cold slice culture dissection medium containing (in mM): 248 sucrose, 26 NaHCO_3_, 10 glucose, 4 KCl, 5 MgCl_2_, 1 CaCl_2_, 2 kynurenic acid and 0.001% phenol red. pH was 7.4, osmolarity 310-320 mOsm kg^-1^, and solution was saturated with 95% O_2_, 5% CO_2_. Tissue was cut into 410 µM thick sections on a tissue chopper and cultured at the medium/air interface on membranes (Millipore PICMORG50) at 37° C in 5% CO_2_. For the first 24 h of incubation, 1 µM (Z)-4-hydroxytamoxifen (Sigma H7904) was added to the slice culture medium to induce Cre-activation. No antibiotics were added, and slice culture medium was partially exchanged (60-70%) twice per week and contained (for 500 ml): 394 ml Minimal Essential Medium (Sigma M7278), 100 ml heat inactivated donor horse serum (Sigma H1138), 1 mM L-glutamine (Gibco 25030-024), 0.01 mg ml^-1^ insulin (Sigma I6634), 1.45 ml 5M NaCl (Sigma S5150), 2 mM MgSO_4_ (Fluka 63126), 1.44 mM CaCl_2_ (Fluka 21114), 0.00125% ascorbic acid (Fluka 11140), 13 mM D-glucose (Fluka 49152). Slice cultures were used for experiments between 12 and 28 days *in vitro*. Mice were housed and bred at the University Medical Center Hamburg-Eppendorf. All procedures were performed in compliance with German law and the guidelines of Directive 2010/63/EU. The study was approved by the local authorities (Amt für Verbraucherschutz, Lebensmittelsicherheit und Veterinärwesen, Hamburg; permission # 42/17).

### Mouse genotyping

Tail biopsies were taken from mice at postnatal day 3-4 and lysed using 75 µl alkali buffer containing (in mM): 25 NaOH, 0.2 Na_2_-EDTA*2H_2_O for 60 min at 95°C and then neutralized using 75 µl 40 mM Tris-HCl. PCR-based genotyping of the Rosa26 locus was performed using the primer combinations 5’-AAGGGAGCTGCAGTGGAGTA-3’ and 5’-CCGAAAATCTGTGGGAAGTC-3’ for WT and 5’-GGCATTAAAGCAGCGTATCC-3’ and 5’-CTGTTCCTGTACGGCATGG-3’ for ChETA *knock-in* alleles and 5’-CCGAAAATCTGTGGGAAGTC-3’ and 5’-ATTGCATCGCATTGTCTGAG-3’ for ArchT *knock-in* alleles.

### Perfusion of animals and preparation of slices

Mice were transcardially perfused with cold phosphate-buffered saline (PBS, pH 7.4), followed by 4% paraformaldehyde in PBS under ketamine (130 mg/kg, WDT) and xylazine (10 mg/kg, WDT) anesthesia. Brains were extracted and post-fixed for 24 h in 4% PFA at 4° C. Before slicing, the tissue was changed to 1x PBS for 20 minutes, then slices of 40 µm were sectioned using Leica vibratome, collected and stored in PBS.

### Immunohistochemistry

Organotypic slice cultures were fixed in 4% paraformaldehyde in PBS for 45 min at room temperature and stored in PBS at 4 ° C until further use. Fixed cultures or brain slices from perfused animals were blocked for 2 hours at room temperature using goat-serum based blocking solution containing (in %): 10 Goat serum (Capricorn), 0.2 Bovine Serum Albumin (Sigma-Aldrich A2153-10G), 0.5 TritonX-100 (Sigma Aldrich T-9284). Primary antibodies were directed against IBA1 (rabbit-anti-iba1 WAKO chemicals 019-19741 1:1000) and tdTomato (mouse-anti-dsRed Takara 632392 1:1000). Secondary antibodies were goat anti-rabbit or anti-mouse conjugated with Alexa dyes 488 and 568 (1:1000), respectively (Life Technologies A11008 and A11004). Antibodies were prepared in carrier solution containing (in %): 1 Goat serum (Capricorn), 0.2 Bovine Serum Albumin (Sigma-Aldrich A2153-10G), 0.3 TritonX-100 (Sigma Aldrich T-9284). Primary antibodies were applied overnight at 4 ° C; before application of secondary antibodies, slices were washed three times 10 min with PBS and then incubated with secondary antibodies for 2 hours at room temperature. Slices were washed for three times with PBS, incubated with DAPI (Molecular Probes, Invitrogen) for 5 min and then mounted (Thermo Scientific 9990402). Slices were imaged using an Olympus FV-1000 confocal microscope and microglia morphology and cell count was analyzed using the Imaris Software (Bitplane).

### Two photon imaging and microglia patch clamp

Organotypic hippocampal slice cultures were placed in the recording chamber of a custom-built two-photon laser scanning microscope based on an Olympus BX51WI microscope with a pE-4000 LED light source for epifluorescence and activation of ChETA or ArchT. Power densities of 470 nm and 520 nm light were measured using a 1918-R power meter (Newport). The microscope was equipped with a WPlan-Apochromat 40× 1.0 NA (Zeiss) or HC Fluotar L 25x 0.95 NA (Leica) objective and controlled by ScanImage 2017b (Vidrio). A Ti:Sapphire laser (Chameleon Ultra, Coherent) controlled by an electro-optic modulator (350-80, Conoptics) was used to image microglia at 1040 nm. Slice cultures were continuously perfused with a HEPES-buffered solution (in mM): 135 NaCl, 2.5 KCl, 10 Na-HEPES, 12.5 D-glucose, 1.25 NaH_2_PO_4_, 2 CaCl_2_, 1 MgCl_2_ (pH 7.4, 308 mOsm) at 27-29° C. Nominally Ca^2+^-free extracellular solution consisted of (in mM): 135 NaCl, 2.5 KCl, 10 Na-HEPES, 12.5 D-glucose, 1.25 NaH_2_PO_4_, 3 MgCl_2_ (pH 7.4). Whole-cell recordings from microglia in stratum radiatum were made with patch pipettes (6–8 MΩ) filled with (in mM): 135 K-gluconate, 4 MgCl_2_, 4 Na_2_-ATP, 0.4 Na-GTP, 10 Na_2_-phosphocreatine, 3 sodium-L-ascorbate, and 10 HEPES (pH 7.2, 295 mOsm kg^-1^). Fluorescence-guided microglia whole-cell patch clamp recordings were performed with a Multiclamp 700B amplifier (Molecular Devices) under Ephus software control [18]. Series resistances were below 20 MΩ. Z-stacks of patch-clamped microglia were acquired to document their morphology. Laser tissue damage was performed in a standardized fashion in a region between three to four microglia at a distance of approximately 35-50 µm by pointing the IR-laser beam to the center of the field of view for several ms.

### Image analysis of microglia baseline surveillance and chemotaxis

#### Microglial baseline surveillance

We combined several analysis steps into a semi-automated workflow. Image data were background subtracted (rolling ball radius = 30 pixels) and median filtered (radius = 1 pixel) with ImageJ. In order to correct for small positional shifts that occurred during acquisition, we registered all images from a time-lapse recording to the first image of the series (Efficient subpixel image registration by cross-correlation, version 1.1, MATLAB Central File Exchange). Images were then binarized using Otsu’s method (threshold set to 50% calculated value). Based on a maximum projection of both images and binary images, a mask was drawn to define the region considered for motility analysis. Within the masked region we counted the number of pixels that remained stable, that were lost, and that were gained throughout the time series. From the covered areas and perimeters, we calculated ramification and surveillance indices.

#### Microglial response to laser-induced tissue damage

We created a semi-automated workflow (Matlab R2016b, MathWorks®, code available on GitHub) to analyze microglial responses to laser-induced tissue damage. Individual images from a time-lapse series (2D, Number, time step) were median filtered (3×3 kernel size), converted to binary form (Otsu’s method), polished using dilation, erosion steps and filling of holes. A spot (eccentricity < 0.5, size > 50 pixel) near the image center was defined as the site at which tissue damage was initiated. The image from the tissue damage time point with detected spot coordinates was presented to the user for visual inspection; the location of the spot was corrected if necessary. For further processing, images prior to the damage time point were averaged (median pixel values) to obtain a background image from which an intensity threshold was calculated (locally adaptive threshold, specificity = 0.5). All images were then background subtracted and binarized using the intensity threshold. The region around the damage center (spot dilated with a 9×9 disc) was masked and objects smaller than 5 pixels were removed. Following polishing using a dilation/erosion (5×5 kernel), step object areas, distances, and positions relative to the damage center sectioned into 36 radial slices were extracted.

## Results and discussion

### ChETA effectively depolarizes microglia in response to blue light illumination

For microglia-specific expression of the light-gated ion channel ChETA, we crossed the tamoxifen-inducible microglia-specific Cre-driver line Cx3cr1^CreERT2^ with the conditional R26-CAG-LSL-ChETA mouse line. To visualize microglia in control experiments, we used the conditional reporter line R26-CAG-LSL-tdTomato (Fig. 1a, b). From those mice and littermate controls lacking ChETA, we prepared organotypic hippocampal slice cultures, a three-dimensional system in which the cellular architecture is well preserved. Application of (Z)-4-hydroxytamoxifen in the culture medium (24 h) activated the cre/loxP-system, leading to selective construct expression in the entire microglia population (Fig. 1c). Although microglia became activated during the cutting process and assumed a morphology similar to pathologically-activated microglia shortly after preparation, they regained their highly ramified resting morphology over 9-12 days *in vitro* (DIV) (Fig. 1d-g). Sholl analysis showed that at 29 DIV, their morphology was indistinguishable from microglia we imaged through cranial windows in intact animals (green markers, Fig. 1h). The fraction of microglia in *stratum radiatum* of organotypic cultures fluctuated between 5-10% of all cells, which is not different from the density we measured *in vivo* (Fig. 1i). Furthermore, microglia membrane properties recorded in mature organotypic slice cultures (membrane resistance, capacitance, resting potential; Fig. 2a, b) were very similar to those reported in acute brain slices [19,20] were not correlated with the age of the culture and not different in cultures prepared from male or female animals (Additional file 1).

**Fig. 1.**
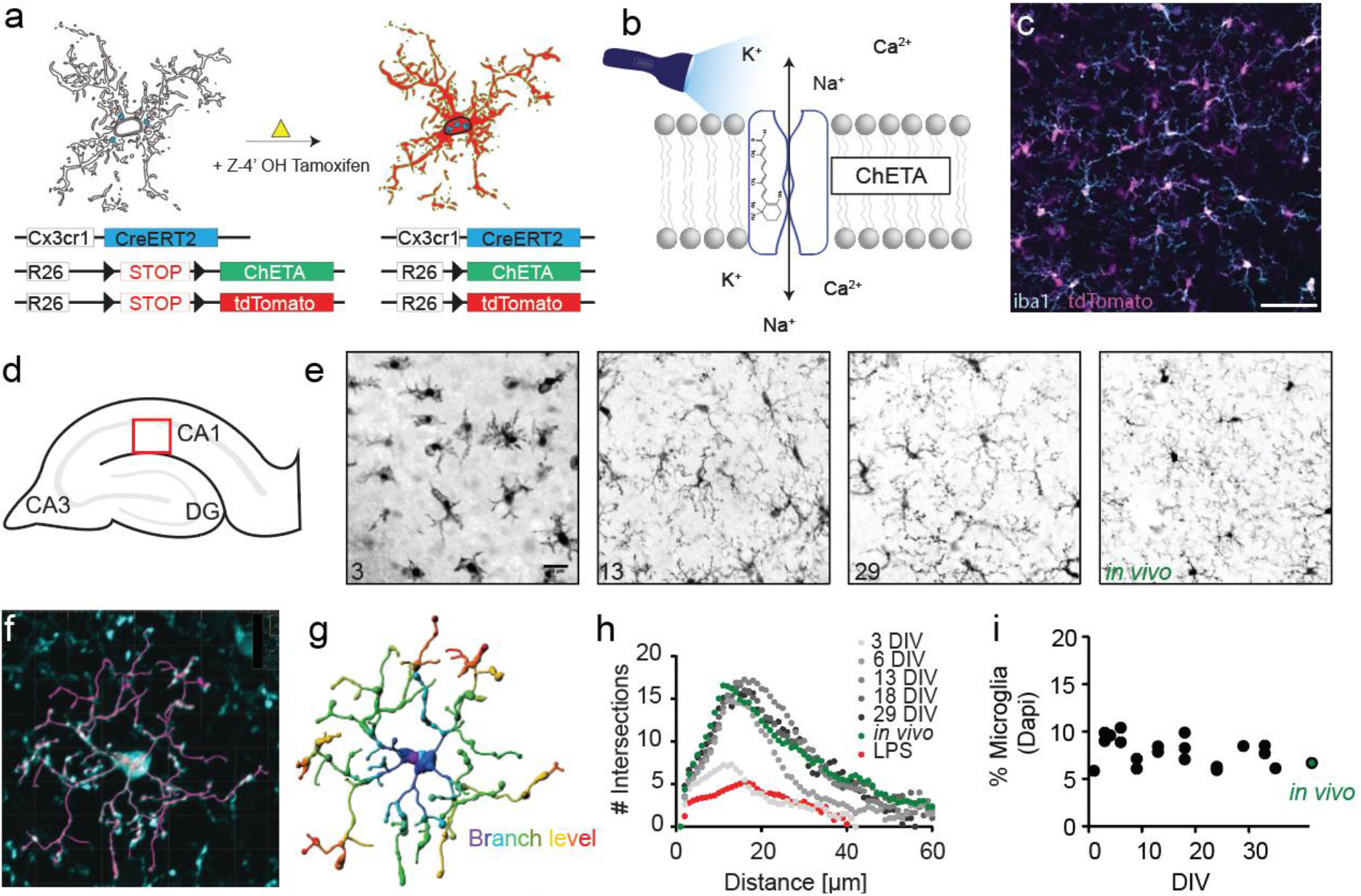
Microglia specific ChETA expression in organotypic hippocampal slice cultures. (a) Schematic overview of microglia-specific expression. The microglia-driver line Cx3cr1-CreERT2 (blue) is crossed with a reporter mouse line (R26-LSL-tdTomato, red) and the ChETA mouse line (R26-LSL-ChETA, green). After injection of (Z)-4-hydroxytamoxifen the tdTomato and ChETA are expressed in microglia (b) Illustration of the Channelrhodopsin-variant ChETA activated by blue light. Scale bar 25 µm (c) Immunostaining using antibodies against the reporter (tdTomato - red) and microglia (iba1 - cyan). (d) Graphic illustration of the hippocampal structure and the investigated area for microglia morphology in E (red square). (e) Z-projection of confocal images acquired for Sholl analysis of microglia at 3, 13, 29 DIV and in vivo. Scale bar: 25 µm. (f) Confocal image of a microglia cell in organotypic slice culture which was fixed with PFA and stained against the microglia marker iba1. Overlay with IMARIS analysis (magenta). (g) Result of microglia branch detection with color coding by branching level. (h) Sholl analysis of microglia over time (number of intersections versus distance from cell body). (i) Quantification of % microglia cells between dentate gyrus and CA1 relative to total cell count (DAPI).

**Fig. 2.**
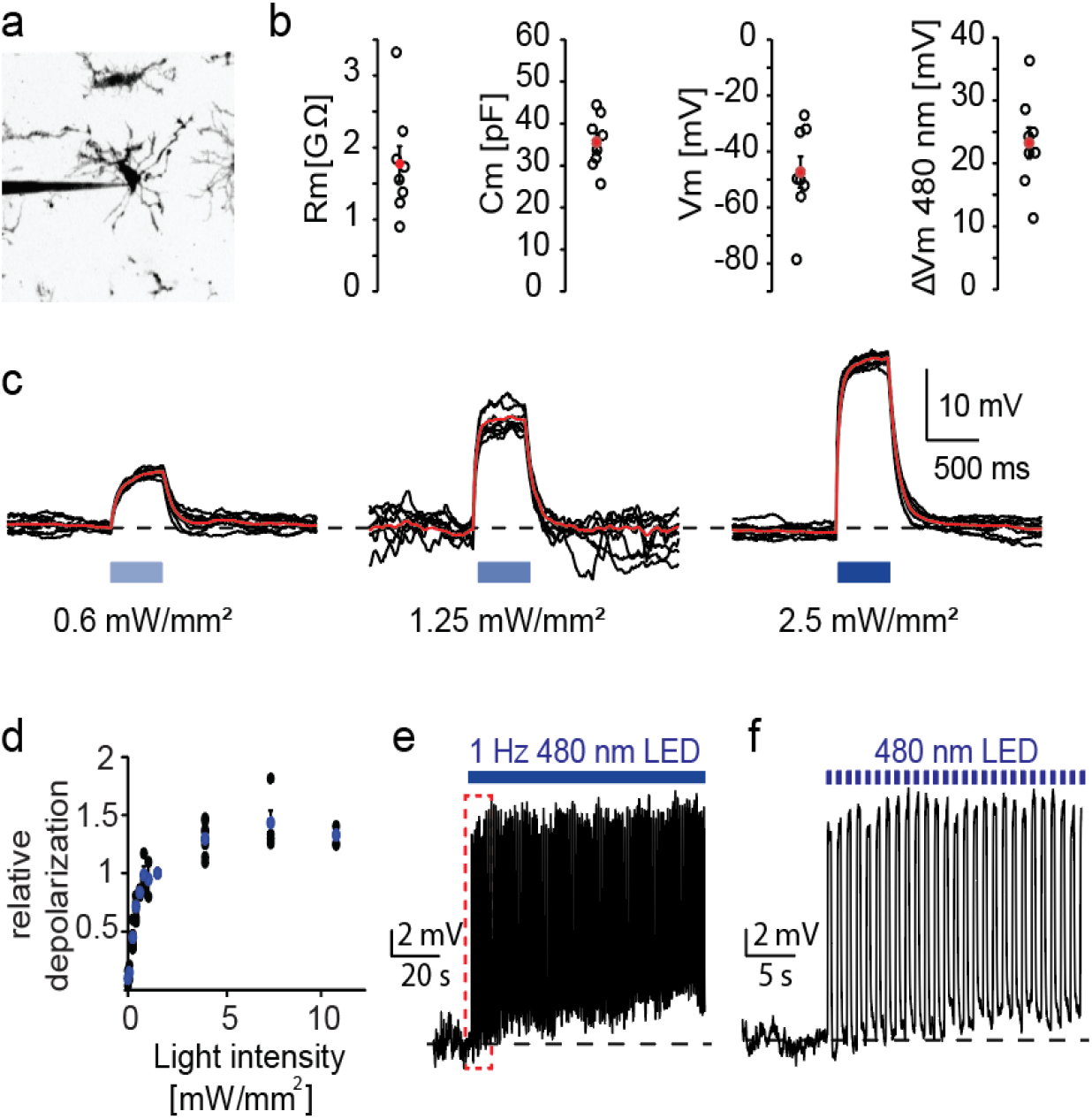
Optogenetic microglia depolarization using the Channelrhodopsin variant ChETA. (a) Two-photon maximum projection of a microglia patched in organotypic slice culture. Color inversion for better visibility. (b) Microglia cell properties; from left to right: Membrane resistance, cell capacitance, membrane potential, change in membrane potential upon illumination with 1 mW/mm^2^ 480 nm light. Black circles: n = 9 microglia, 5 slices; red dot: mean ± SEM. (c) Light-induced membrane depolarizations in microglia. Black: individual sweeps (n = 17 sweeps), red: peristimulus time histogram (PSTH) of seven consecutive sweeps with the same light intensity. (d) Relative depolarization of microglia membrane potential 480 nm light. Blue dots: mean ± SEM. (e) Voltage-clamp recording of a microglia cell with repetitive 1 Hz 480 nm light stimulation at 1 mW/mm^2^ (n = 7 microglia, 3 slices, female, DIV 12-20) (f) Enlargement of red dotted box in e.

The channelrhodopsin-2 variant ChETA (ChR2/E123T/H134R) was originally developed for fast response kinetics at the expense of smaller photocurrents [21]. Therefore, relatively high light intensities and expression levels are required to spike neurons [22]. Microglia, however, exhibit an almost 10-times higher input resistance compared to neurons, so small photocurrents generate large changes in microglia membrane potential [4]. Using increasing blue light intensities, we achieved a maximal depolarization of ΔVm = 36 mV from the resting potential of microglia (Fig. 2b, d). During high intensity light pulses, depolarization reached a plateau at 0 mV, the reversal potential of channelrhodopsin [22]. Even prolonged repetitive depolarization using light flashes at 1 Hz frequency did not lead to desensitization, but produced photocurrents of constant amplitude (Fig. 2e, f). In summary, our genetic approach guaranteed specific expression of the channelrhodopsin variant ChETA exclusively in microglia and allowed for rapid light-controlled depolarization in a dose-dependent and cell-type specific fashion.

### Optogenetic depolarization of microglial membrane potential mildly affects microglia morphology and baseline surveillance

Resting microglia constantly monitor their surrounding parenchyma and the surveyed area greatly depends on their morphology. In the resting state, microglia scan larger areas due to the higher number and greater length of processes compared to pathologically activated microglia [23,24]. The morphological change during pathological activation is often accompanied by strong changes in microglia membrane potential [25–28]. Thus, a direct link between morphology and membrane potential is plausible. Consistent with this, Madry *et al*. found that pharmacological block of the constitutively open outward-rectifying THIK-1 potassium channel, or genetic knock-out thereof, leads to a more depolarized membrane potential in microglia and a more activated morphological phenotype with reduced surveillance [2]. To investigate whether microglia depolarization by itself causes motility changes, we optically modulated the membrane potential (480 nm light flashes for 20 min) while tracking microglia motility over 50 minutes with two-photon microscopy. Optogenetic depolarization did not induce any morphological changes and had only minor, transient effects on baseline motility compared to microglia in control slices that did not express ChETA (Additional file 2). Taken together, our data show that strong short-term depolarization of microglia membrane potential does not induce any lasting changes in morphology or surveillance behavior.

### Optogenetic microglia depolarization decelerates chemotactic response kinetics

Microglia maintain a negative resting potential around -40 mV and rapidly hyperpolarize further when sensing extracellular ATP or other nucleotides. This instantaneous hyperpolarization is mediated via the activation of P2Y12 receptors, which results in opening of THIK-1 channels and occurs simultaneously with the onset of microglia process movement towards the source of ATP [2,14]. Hyperpolarization could be part of the trigger mechanism for the rapid chemotactic response, or just an epiphenomenon. To find out, we altered microglia membrane potential during the chemotactic response using optogenetic depolarization.

After recording microglia baseline motility for 15 minutes with two-photon microscopy, we induced spatially precise tissue damage in a region surrounded by three or four microglia by parking the IR-laser in the center of the field of view for 1.5 seconds. The distance between the tips of microglia processes and the laser focus was 35-50 µm (Fig. 3a, b). Microglia rapidly responded to the laser damage by extending their processes and within a few minutes, the site of damage was completely surrounded by pseudopodia.

**Fig. 3.**
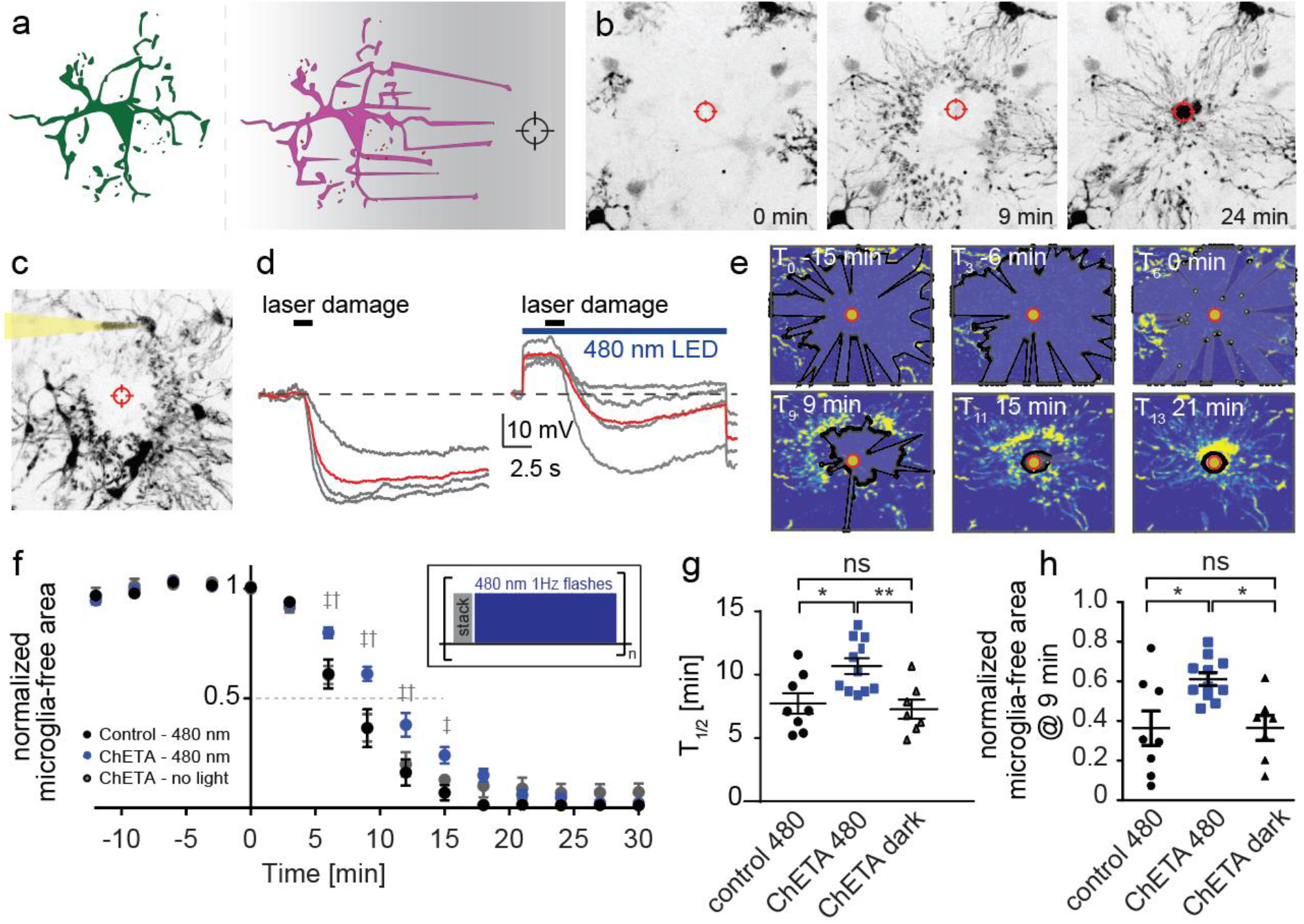
Optogenetic microglia depolarization decelerates chemotactic response kinetics. (a) Graphic illustration and representative images of microglia chemotaxis towards an induced laser-damage. (b) Two-photon maximum projections of the chemotactic response 0, 9, 24 min after the laser-damage. (c) Two-photon z-projection of a patched microglia during chemotaxis. (d) Voltage-clamp recordings of patched microglia during chemotaxis. Gray: Individual microglial responses from four experiments, red: Average of all experiments. Left: no light stimulation during laser-damage, right: with light stimulation during laser damage. (e) Automated MATLAB analysis of chemotaxis quantified as the reduction in microglia-free area around the laser damage (black polygon) at different time points of the experiment. (f) Relative laser damage response measured as microglia-free area. Black: Control slices (no construct) with light stimulation (n=8 areas, 5 slices). Gray: Experiments with ChETA expression in microglia, but without light stimulation (n=7 areas, 4 slices). Blue: Slices with ChETA expression in microglia combined light stimulation (n = 11 areas, 7 slices). Insert: Graphic representation of light stimulation protocol between stack acquisitions. 2-way ANOVA (‡ control480 – ChETA480 p<0.001, († ChETA no light – ChETA480 p<0.01) (g) Time to 50% engulfment was prolonged by optogenetic depolarization. (h) 9 min after injury, the microglia-free area was larger when microglia were depolarized. G and H: One-way ANOVA with Tukey’s post-hoc comparison (*p < 0.05, **p<0.01, ***p<0.001)

To investigate how effectively optogenetic depolarization reduced ATP-mediated hyperpolarizing currents we performed whole-cell patch clamp recordings from microglia during laser damage (Fig. 3c, d). Using 480 nm illumination before and during tissue damage, we were able to substantially reduce the amount of hyperpolarization in microglia (Fig. 3d). In addition to light stimulation during laser damage, we illuminated the field of view with 1 Hz flashes at 480 nm between image acquisition to depolarize microglia throughout the chemotactic response (Fig. 3f, insert).

After determining the light intensities and frequencies required to effectively depolarize microglia, the actual chemotactic experiments were then performed without patch-clamp electrodes. We included two types of control experiments: Microglia expressing only the fluorescent protein (no ChETA) and microglia that expressed ChETA, but were not illuminated. These control groups showed similar response kinetics in response to laser damage, demonstrating that neither ChETA expression nor light application alone had an effect on microglia responsiveness (T½ _ctrl_ = 7.7 ± 0.7 min, free area_ctrl_ = 36% ± 8%; T½ _ChETA_nolight_ = 7.3 ± 0.7 min, free area_ChETA_nolight_ = 37% ± 6%; Fig. 3g, h). In contrast, microglia expressing ChETA exhibited significantly slower response kinetics when illuminated compared to both control microglia with light and ChETA expressing microglia without light stimulation (T½ _ChETA+light_ = 10.7 ± 0.6 min, free area_ChETA+light_ = 61% ± 3%, Fig. 3G, H). Microglia response kinetics were not correlated with distance from the laser focus, temperature, days in vitro, or sex (Additional file 3).

To test the effect of artificial hyperpolarization, we expressed the light-driven proton pump ArchT in microglia. Illumination with yellow light produced reliable hyperpolarization of about -40 mV (Additional file 4). When we combined ArchT-mediated hyperpolarization with laser injury, the result was not additive as in ChETA experiments, but the electrical response to injury was absent or even reversed. Consequently, chemotaxis was not different in illuminated vs non-illuminated microglia (Additional file 5). These results are in line with a tight coupling between membrane potential and speed of pseudopodia extension. As ArchT is known to alter intracellular pH, however, effects on ion currents are difficult to interpret in detail.

In microglia, an increased frequency of Ca^2+^ transients is correlated with pathophysiological activation and chemotactic responses. Cellular processes moving towards a source of extracellular ATP exhibit significantly more Ca^2+^ transients compared to resting microglia or processes that are located away from the tissue damage [11], and pseudopodia could be actively guided towards the ATP source by local Ca^2+^ influx. Thus, it is plausible that optogenetic depolarization slows down chemotaxis by reducing the driving force for Ca^2+^. To investigate whether response kinetics depend on Ca^2+^ entry, we performed a series of chemotactic experiments in nominally Ca^2+^-free extracellular solution. In Ca^2+^-free extracellular solution, process extension towards the damaged area was significantly slowed down (ΔT½ = 3.1 min; Additional file 6), very similar to the slowing induced by optogenetic depolarization (ΔT½ = 3.0 min).

Together, these data demonstrate that microglia hyperpolarization is an important component of the rapid chemotactic response and suggest that the coupling of membrane potential to process elongation might be mediated by enhanced influx of Ca^2+^ ions. However, the onset of chemotactic responses was not affected, neither in optogenetic depolarization nor in Ca^2+^-free extracellular solution, indicating that this important cellular function is controlled by multiple pathways that act in parallel, partially redundant fashion. Some of them, e.g. G-protein coupled P2Y_12_ receptors are not at all expected to depend on membrane potential or calcium influx. Thus, controlling the membrane potential of microglia can significantly slow down, but not completely switch off the injury response.

### Microglia hyperpolarization is not an epiphenomenon

Microglia are the only resident immune cells in the brain and although they maintain a negative resting potential and rapidly respond to an external stimulus (ATP) with strong transient hyperpolarization, they are considered non-excitable cells. Yet, microglia exhibit reliable changes in membrane potential upon external stimulation, which in turn affect crucial microglia functions such as chemotaxis or phagocytosis [29,30]. Here we provide evidence that membrane hyperpolarization is not just a byproduct, but part of the signaling pathway for rapid stimulus-response coupling. Previous studies investigating the depolarizing effect of pharmacological block of K^+^ channels on chemotactic response kinetics yielded conflicting results: While K^+^ channel block with quinine reportedly abolishes chemotaxis, K^+^ channel block with tetrapentylammonium (TPA) did not seem to affect response kinetics [2,29]. It is difficult to determine whether the stimulation strength was comparable in both studies, as chemotaxis was induced by pressure application of ATP at different concentrations and evaluated with different metrics. In both studies, chemotaxis assays were performed in acute brain slices which had sustained tissue damage during the slicing process. The state of microglia activation at baseline might therefore depend on subtle details of the slice-making and incubation procedure, leading to poor reproducibility across labs. Microglia in mature organotypic cultures may provide a more stable baseline for chemotaxis and other assays. In addition, our optogenetic approach reduces the risk of unspecific side effects that is the bane of pharmacology. Madry et al. also generated THIK-1 knock-out mice in which microglia are chronically depolarized and show no ATP-induced currents. The chemotactic response was not tested in these animals, but THIK-1-KO microglia had fewer and less motile processes at baseline, suggesting microglia activation. As we show here, acute optogenetic depolarization does not change the morphology and hardly affects surveillance activity of microglia. Thus, it is likely that chronic depolarization triggers changes in expression profile, e.g. downregulation of P2Y12 and upregulation of A2A receptor expression [31]. While P2Y12 receptors are responsible for the chemotactic response, A2A receptors mediate process retraction from ATP in activated microglia [2,31].

### Hyperpolarization increases the driving force for calcium ions

Ca^2+^ signaling is known to play a key role in microglia activation. The main entry route is not via voltage-gated Ca^2+^ channels, which are only expressed under pathological conditions [32– 34], but through ligand-gated receptors with high Ca^2+^ permeability. Thus, in contrast to neurons, depolarization reduces Ca^2+^ influx in microglia. A direct link between hyperpolarization and Ca^2+^ signaling has been demonstrated in many types of immune cells and is strengthened by the finding that under resting conditions, microglia exhibit almost no Ca^2+^ transients (4% of resting microglia show a spontaneous Ca^2+^ transient during a 20 min recording session [11]). Pathological hyperpolarization strongly affects microglia function and is accompanied by an increased number and intensity of Ca^2+^ transients [35,36]. In particular, potassium channel expression plays an important role in the regulation of the hyperpolarized membrane potential (for review see [37–39]). In addition, it was shown that chemotactic responses are accompanied by local Ca^2+^ transients in processes approaching the source of ATP. Optogenetic depolarization might effectively decrease Ca^2+^ currents through ligand-gated receptors, thereby decelerating chemotactic responses. Corroborating evidence comes from our chemotaxis experiments in Ca^2+^-free saline which phenocopied the effect of optogenetic depolarization. Actin polymerization, one of the major driving forces for pseudopod extension during chemotaxis, is highly regulated by Ca^2+^-activated enzymes. In our optogenetic depolarization experiments, we were only able to slow down process extension and not completely prevent all movements, which might be due to our pulsed light protocol in combination with a fast closing opsin. However, as the reversal potential for Ca^2+^ is far above the reversal potential of ChETA (−5 mV), even constant illumination would not prevent Ca^2+^ influx completely. The remaining slowed down chemotactic response may be driven by residual calcium influx or by parallel, Ca^2+^-independent signaling pathways.

## Conclusions

We have shown that ATP-mediated hyperpolarization is not just an epiphenomenon of microglia activation, but part of the signal transduction process controlling the rapid extension of pseudopodia towards the site of injury. Microglia could therefore be considered excitable cells, but they react in an inverted fashion when compared to neurons: Depolarization dampens their reactivity while hyperpolarization is part of their active response to extracellular danger signals. Beyond chemotaxis, optogenetic control of microglia membrane potential opens the possibility to investigate other physiological processes that may depend on membrane potential, such as cytokine release via exosomes, without affecting other cell types.

## Abbreviations

ACSF: Artificial cerebrospinal fluid
CAG: strong synthetic promoter
ChETA: Channelrhodopsin E to T, accelerated
CX3CR1: CX3C chemokine receptor 1
IBA1: ionized calcium-binding adapter molecule 1
Kir2.1: voltage-gated potassium channel, inward rectifier
Kv1.3: voltage-gated potassium channel, delayed rectifier
LSL: Lox-STOP-lox cassette
P2X4: purinergic receptor, ligand-gated ion channel
P2X7: purinergic receptor, ligand-gated ion channel
P2Y12: purinergic G_i_-protein coupled receptor
PBS: phosphate-buffered saline
R26: ROSA26 locus for ubiquitous gene expression
tdTomato: dimeric red fluorescent protein
THIK-1: tandem-pore domain halothane-inhibited K^+^ channel 1
TRP: transient receptor potential channel

## Declarations

### Ethics approval and consent to participate

Animal experiments were approved by the local authorities (Amt für Verbraucherschutz, Lebensmittelsicherheit und Veterinärwesen, Hamburg, Germany; Permission # 42/17).

### Consent for publication

Not applicable

### Availability of data and materials

The datasets used and/or analyzed during the current study are available from the corresponding authors on reasonable request.

### Author’s contributions

LL and TGO designed the experiments and prepared the manuscript. LL performed microglia patch clamp recordings, microglia chemotaxis and motility experiments, immunohistochemical experiments and all data analysis. MLB performed microglia motility experiments and immunohistochemical experiments. CS wrote software to analyze microglia motility and chemotaxis. All authors read and approved the final manuscript.

## Acknowledgements

We thank I. Ohmert and S. Graf for the preparation of organotypic cultures and excellent technical assistance. We thank M. Fuhrmann for providing us with two male mice of the Jackson Laboratory (JAX) mouse lines JAX 020940 and JAX 007909. We thank I. Hanganu-Opatz for providing us with a breeding pair of the JAX mouse lines 017455 and 021188.

## Supplementary Information

**Additional file 1.**
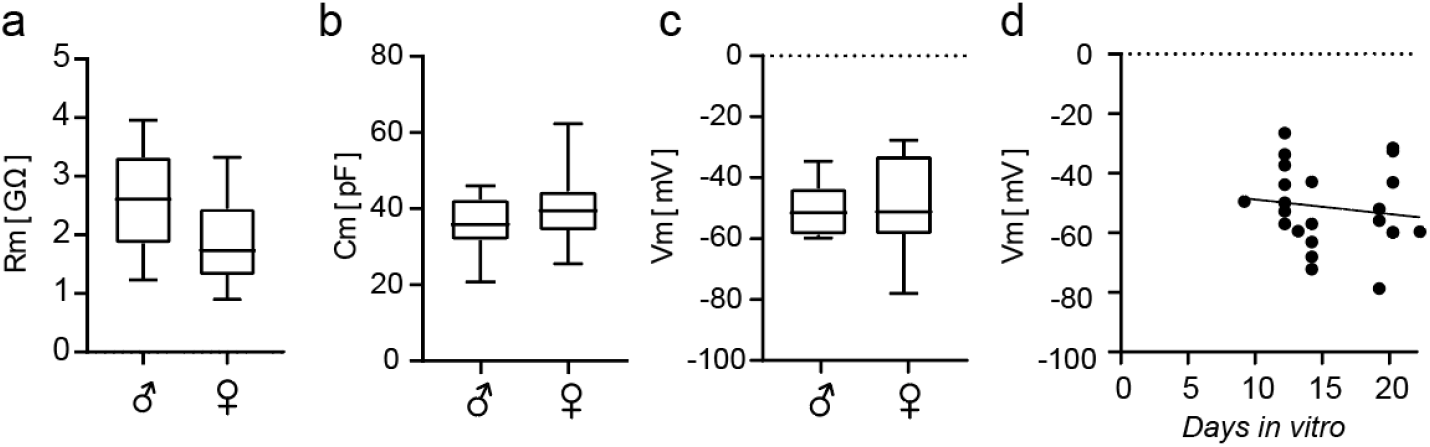
Cell parameters in microglia of male and female mice show no differences in slice cultures. Duration of slice culture in vitro has no significant effect on microglia membrane potential. (a) Membrane resistance (n=14/9, male/female), (b) Membrane capacitance (n=9/12, male/female), (c) Membrane resting potential (n=12/8, male/female), (d) Membrane resting potential vs. days in vitro (n=23). Spearman’s ρ = -0.24, P = 0.27, n = 23 experiments.

**Additional file 2.**
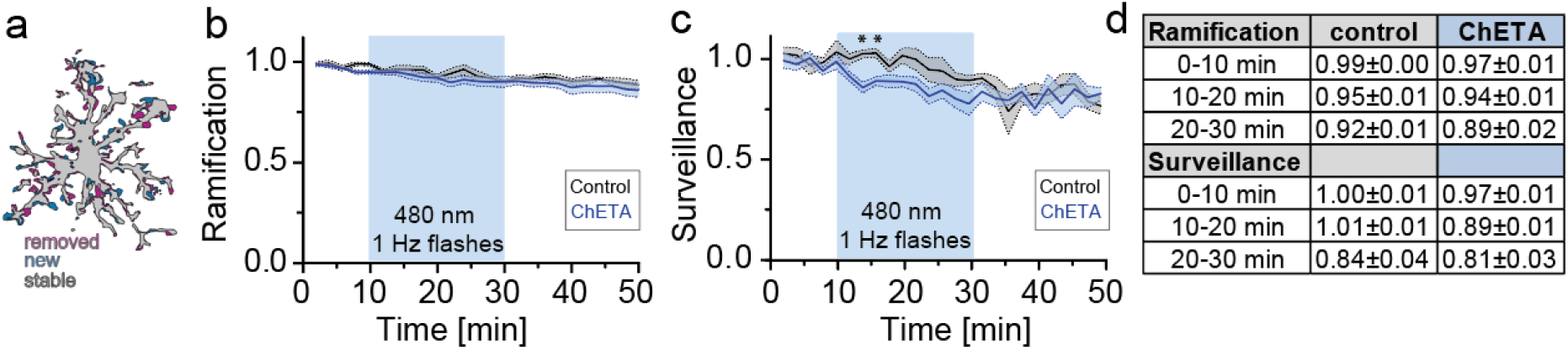
Optogenetic depolarization mildly affects microglia baseline properties. (A) Superimposed images of the same microglia at two timepoints one minute apart showing process movement during surveillance (purple, retracted; blue, extended and gray stable processes). (B-C) Relative microglia ramification (B) and surveillance index (C) over the time course of 50 min (ChETA_480, n = 4 microglia in 3 slices, control_480, n = 8 microglia in 5 slices). Light application (20 min with 480 nm) was started after 10 min of baseline recording. Statistical analysis: Two-way ANOVA with Bonferroni post-hoc comparison (*p < 0.05) (D) Overview table of relative ramification and surveillance indices in the 10 min before light application and the first 10 minutes of light application (Average ± SEM).

**Additional file 3.**
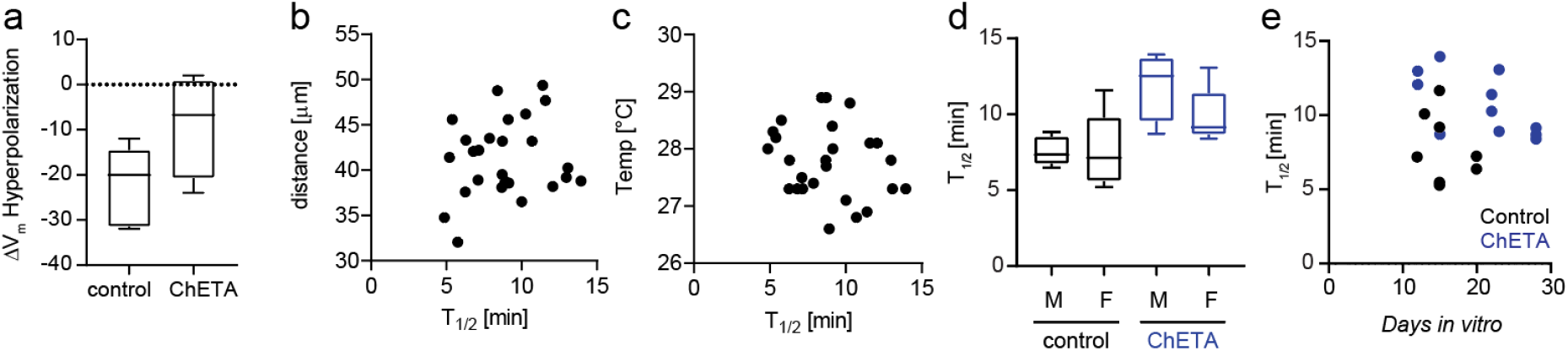
Microglia response time is independent from distance of ablation, temperature, sex, or days *in vitro*. (a) Amount of hyperpolarization of microglia membrane potential induced by laser-damage with and without activation of ChETA (n=5 controls, n=4 ChETA, DIV 20-22) (b) T_1/2_ plotted against the distance of microglia processes from the laser damage (Spearman’s ρ = 0.15, P = 0.47, n = 26 experiments). (c) T_1/2_ plotted against the temperature of the extracellular solution (Spearman’s ρ = −0.24, P = 0.25, n = 26 experiments). (d) T_1/2_ plotted for microglia in slice cultures from male (n=5/4 control/light) and female (n=8/7 control/light) animals. (e) T_1/2_ plotted against DIV for control slices (black) and ChETA slices (blue). Control: Spearman’s ρ = -0.2, P = 0.66, n = 8 experiments, ChETA: Spearman’s ρ = -0.6, P = 0.06, n = 11 experiments.

**Additional file 4.**
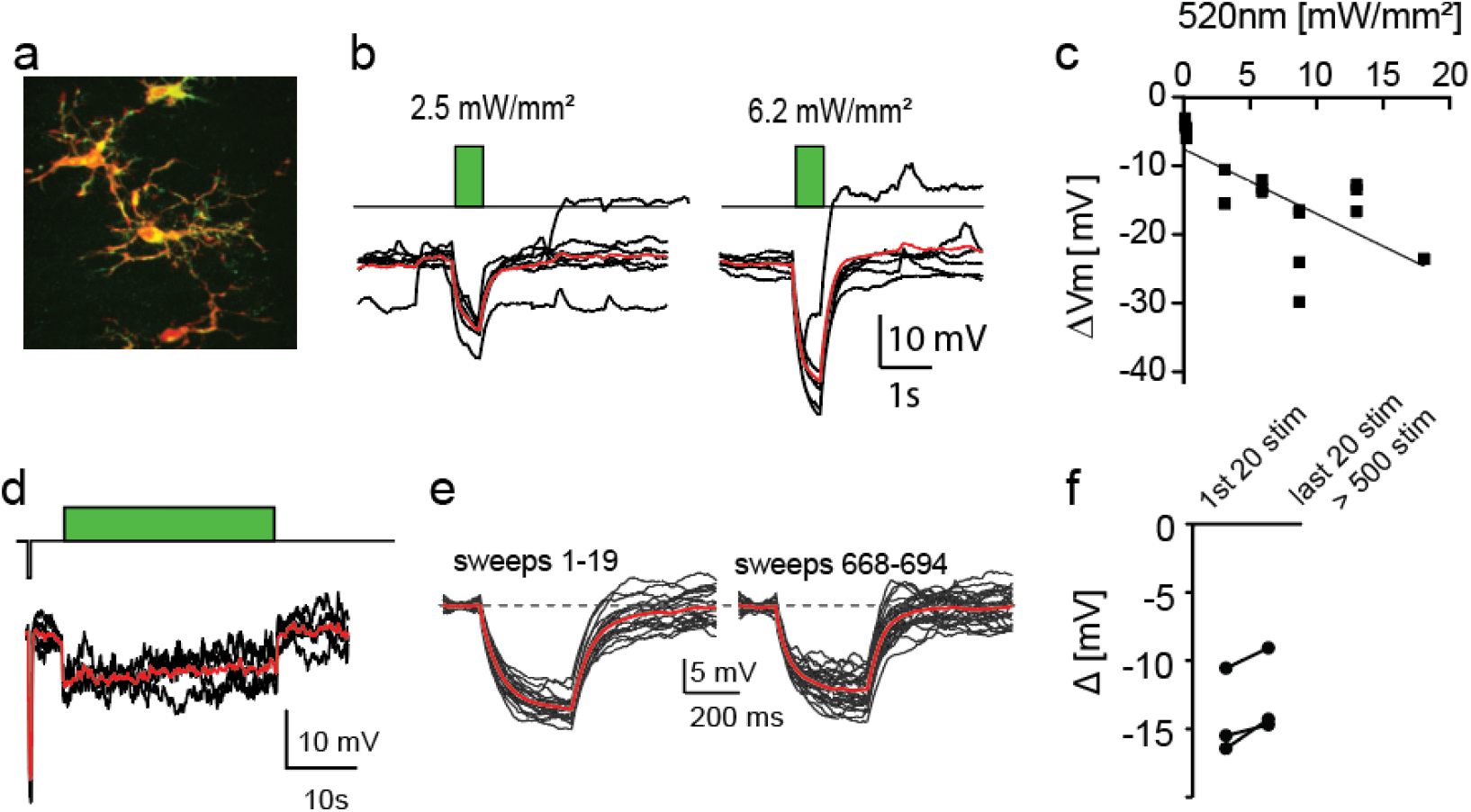
ArchT effectively hyperpolarizes microglia under resting conditions. (a) Immunohistochemical staining of ArchT (green) in microglia (red). (b) Light -induced currents in microglia expressing ArchT. Red lines: Average time course. (c) Light-dose dependence of currents induced in microglia. (d) Prolonged light stimulation induces robust, lasting currents in microglia. (e-f) Repetitive light stimulation induces stable light-induced currents in microglia.

**Additional file 5.**
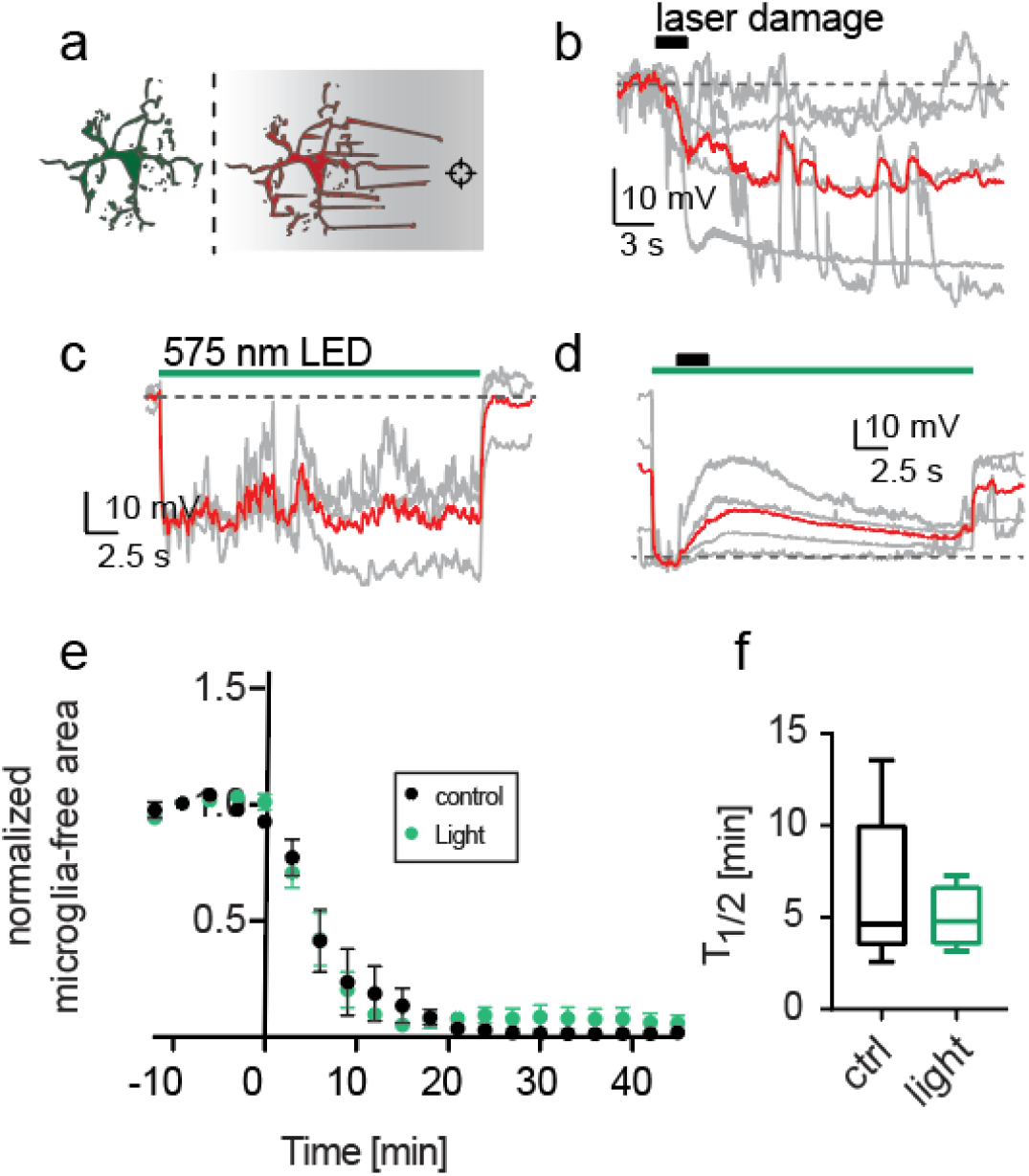
ArchT activation during laser damage does not further increase hyperpolarization and has no effect on laser-damage induced chemotactic responses. (a) Schematic drawing of a chemotactic microglia response. (b) ATP-mediated hyperpolarization in microglia. Red line: Average time course. (c) ArchT light-induced hyperpolarization. (d) Combination of ArchT induced hyperpolarization with laser-damage response. (e) Chemotactic response over time while constantly applying light during acquisition of stacks. (n = 9 (5 female, 4 male)) (f) T1/2 of microglia responses in control and ArchT expressing cultures.

**Additional file 6.**
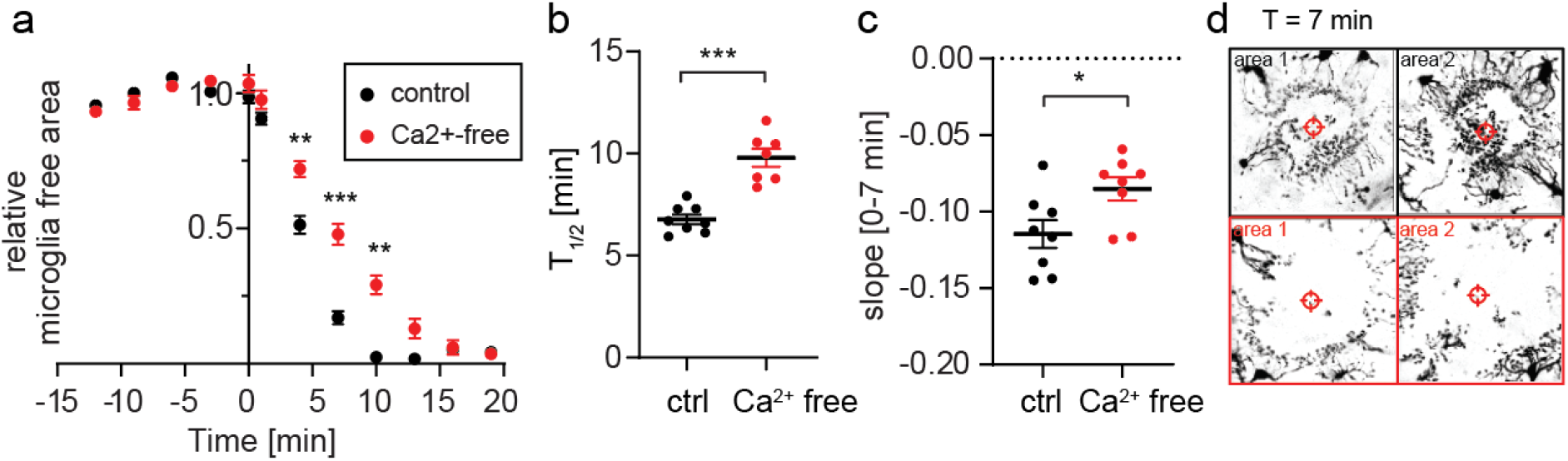
Microglia response kinetics towards tissue damage are slowed down in nominally Ca^2+^ free extracellular solution. (a) Relative laser damage response measured as microglia-free area in HEPES solution with 2 mM Ca^2+^ (n = 8, 3 slices, female, DIV 15-21) (black) and in nominally Ca^2+^-free HEPES solution (n = 8, 3 slices, female, DIV 15-16) (red). 2-way ANOVA (** p < 0.01, *** p < 0.001). (b-c) Summary of T_1/2_ and microglia-free area slope (0-7 min). One-way ANOVA with Tukey’s post-hoc comparison (* p < 0.05, *** p < 0.001). (d) Z-projection of individual areas 7 min after laser damage. Black: microglia processes extension in control conditions with 2 mM Ca^2+^. Red: microglia process extension under nominally Ca^2+^-free conditions.

